# Occurrence of plant-specific 4/1 gene in streptophyte algae: brief view on the gene and protein evolution

**DOI:** 10.1101/2023.12.28.573512

**Authors:** Sergey Y. Morozov, Andrey G. Solovyev

## Abstract

Previously, the plant-specific 4/1 proteins have been found to be encoded by single-copy genes in most land plants (clade *Embryophyta*) but not in green algae. We first identified and characterized 4/1 genes in arabidopsis (At-4/1) and tobacco (Nt-4/1). Importantly, the 4/1 proteins in *Magnoliophyta* species are characterized by a highly conserved C-terminal domain of 30-37 amino acids. In this paper, we report the analysis of 4/1 genes in streptophyte algae – ancestors of lower land plants. AUGUSTUS *ab initio* gene prediction was used to predict 4/1 protein-coding genes in the chromosomal DNA sequences of several algae from classes *Mesostigmatophyceae, Klebsormidiophyceae* and *Zygnemophyceae*. Interestingly, in contrast to an inronless 4/1-like gene previously found in another charophyte alga *Chara braunii*, these genes contain several introns that is consistent with the 4/1 exon-intron organization of land plants. In general, the chromosomes of the studied charophyte algae were found to encode Magnoliophyta-like 4/1 proteins that share their previously described general gene structure and protein properties. These new data on the 4/1-like genes and proteins in the Streptophyta clade suggest that 4/1 proteins are probably function as accessory factors in stress response, but these polypeptides are not required for the primary metabolic functions of streptophyte cells.

## 1. Introduction

The 4/1 protein (At-4/1) was first identified among the *Arabidopsis thaliana* polypeptides interacting with the *tomato spotted wilt tospovirus* movement protein (MP) [1] and capable of MP-independent intracellular movement that gives rise to the 4/1-containing bodies located close to the cell wall and, presumably, plasmodesmata [2,3]. Moreover, the 4/1 protein of *Nicotiana tabacum* (Nt-4/1) has the ability of nucleo-cytoplasmic trafficking. Nuclear export of 4/1 requires the activity of CRM1/exportin 1, whereas transport of the protein from the cytoplasm to the nucleus is dependent on the bipartite nuclear localization signal (NLS) [3]. In addition, some cytoplasmic 4/1-containing bodies are found to align with actin microfilaments and are capable of actin-dependent movement [2-5].

The exact functions of 4/1 proteins in plants are still unclear. However, the data available in the NCBI gene expression omnibus (GEO) show that arabidopsis and rice 4/1 mRNA levels often increase in response to various biotic and abiotic stresses, such as anoxia and fungal infection. This suggests that 4/1 expression may be controlled to maintain plant homeostasis [3,4]. In addition, we have shown that virus-induced gene silencing of the *Nicotiana benthamiana* single-copy 4/1 gene (Nb-4/1) resulted in weakened plant control of viroid pathogen spread [3,6,7], and that the Nt-4/1 protein responds rapidly to mechanical and temperature stresses by relocalizing into numerous small bodies that are likely associated with the cortical ER and containing increased levels of the putative 4/1 interactor protein VAP27 [8]. Importantly, a recent paper shows that VAP27 molecules serve to recruit other polypeptides or protein complexes to the ER–plasma membrane contacts containing plant mechanosensitive ion channel MSL10 [9].

Regarding the possible functions of the 4/1 proteins, it is important to note that during the ontogenesis of *N. benthamiana* plants, the activity of the Nt-4/1 promoter is first detected in cotyledons in association with veins. At later developmental stages, the promoter activity is detected in association with veins of younger leaves and stem internodes, with the hypocotyl being the most intensively expressing part of developing plants. The Nt-4/1 promoter activity is cell type-specific. It is mostly restricted to xylem parenchyma and phloem parenchyma in stems, while in young or small leaf veins it is detected in primary phloem cells [7].

The 4/1 proteins are mostly alpha-helical polypeptides containing up to five-six coiled-coil motifs [4,6]. Our studies suggest that Nt-4/1 consists of three structural domains that have independent structural folds [6]. The central and C-terminal domains have been shown to efficiently bind imperfect double-stranded RNAs including viroid RNA. Importantly, the most C-terminal coiled-coil motif in the Nt-4/1 protein, which represents a highly evolutionarily conserved region of land plant 4/1 proteins, contains a high affinity RNA binding site [4,6,10].

Most 4/1 genes encoded by land plant genomes of have been shown to contain eight exons and seven introns with some exceptions [2,3,6,10]. Several species of streptophyte algae from the classes *Zygnematophyceae* and *Coleochaetophyceae* (e.g., *Spirogyra pratensis* and *Chaetosphaeridium globosum*) also appear to encode 4/1-like proteins, as predicted from transcriptomic analyses. However, the exon-intron organization of the corresponding algal genes remained unknown [10]. Initially, a nearly complete genomic sequence was reported for another streptophyte alga *Chara braunii* (class *Charophyceae*) [11], and we predicted that this genome contains a 4/1-like locus that is intronless [12]. Accordingly, we have hypothesized that the intronless *Chara* 4/1-like gene is initially originated by retrotransposon-dependent gene transfer to the algal genome and then subjected to new intronization steps in early land plants [12].

Modern phylogenetic tree reconstructions of the Viridiplantae based on nuclear and plastid transcriptome/genome sequences have shown that charophycean alga of the classes *Zygnematophyceae* and *Coleochaetophyceae* are the first and second closest algal relatives of land plants, respectively [13-15]. In this paper, recent studies on chromosomal DNA sequences of several streptophyte algae from the basal classes *Mesostigmatophyceae* and *Klebsormidiophyceae* together with the data on *Zygnemophyceae* genomes [13,14] allow us to verify our proposals on the origin of 4/1-like genes in *Charophyceae* and to highlight pathways in the evolution of the land plant 4/1 gene organization.

## 2. Results

In last years, several studies have been published on the genome sequences of charophyte algae. These algae include the basal species *Chlorokybus* atmophyticus (family *[Chlorokybaceae](https://www.ncbi.nlm.nih.gov/Taxonomy/Browser/wwwtax.cgi?mode=Undef&id=131213&lvl=3&lin=f&keep=1&srchmode=1&unlock)*; class *[Chlorokybophyceae](https://www.ncbi.nlm.nih.gov/Taxonomy/Browser/wwwtax.cgi?mode=Undef&=131213&lvl=3&lin=f&keep=1&srchmode=1&unlock)*) and *Mesostigma viride* (family *[Mesostigmataceae](https://www.ncbi.nlm.nih.gov/Taxonomy/Browser/wwwtax.cgi?mode=Undef&id=35428&lvl=3&lin=f&keep=1&srchmode=1&unlock)*; class *[Mesostigmatophyceae](https://www.ncbi.nlm.nih.gov/Taxonomy/Browser/wwwtax.cgi?mode=Undef&id=96475&lvl=3&lin=f&keep=1&srchmode=1&unlock)*) [16] as well as *Klebsormidium nitens (family [Klebsormidiaceae](https://www.ncbi.nlm.nih.gov/Taxonomy/Browser/wwwtax.cgi?mode=Undef&id=3173&lvl=3&lin=f&keep=1&srchmode=1&unlock);* class *[Klebsormidiophyceae](https://www.ncbi.nlm.nih.gov/Taxonomy/Browser/wwwtax.cgi?mode=Undef&id=131220&lvl=3&lin=f&keep=1&srchmode=1&unlock))* [17] and *Chara braunii (*family *[Characeae](https://www.ncbi.nlm.nih.gov/Taxonomy/Browser/wwwtax.cgi?mode=Undef&id=3146&lvl=3&lin=f&keep=1&srchmode=1&unlock);* class *[Charophyceae](https://www.ncbi.nlm.nih.gov/Taxonomy/Browser/wwwtax.cgi?mode=Undef&id=304574&lvl=3&lin=f&keep=1&srchmode=1&unlock)*) [11]. In addition, genomes were sequenced for a dozen species and strains of the class *Zygnematophyceae* including *Mesotaenium endlicherianum (*family *[>Mesotaeniaceae](https://www.ncbi.nlm.nih.gov/Taxonomy/Browser/wwwtax.cgi?mode=Undef&id=31319&lvl=3&lin=f&keep=1&srchmode=1&unlock)*; order *Zygnematales*) [18], *Penium margaritaceum* (family *Peniaceae*; order *Desmidiales*) [19] and some Closterium species (family *[Closteriaceae](https://www.ncbi.nlm.nih.gov/Taxonomy/Browser/wwwtax.cgi?mode=Undef&id=131211&lvl=3&lin=f&keep=1&srchmode=1&unlock)*; order *[Desmidiales](https://www.ncbi.nlm.nih.gov/Taxonomy/Browser/wwwtax.cgi?mode=Undef&id=131210&lvl=3&lin=f&keep=1&srchmode=1&unlock)*) [20].

### 2.1. Search for 4/1 protein conserved signatures encoded by charophyte algae

To reveal potential 4/1 loci candidates in charophyte algae genomes, we performed an initial TBLAST search in the NCBI WGS and TSA databases using as a query the concatenated dataset of conserved C-terminal signatures in the 48 land plant-encoded 4/1 proteins (see Table 3 in [10]). Significant protein conservation in the C-terminal regions was revealed for the protein-encoding regions of chromosomal loci in six algal species and cDNAs in three species (Figure 1).

**Figure 1.**
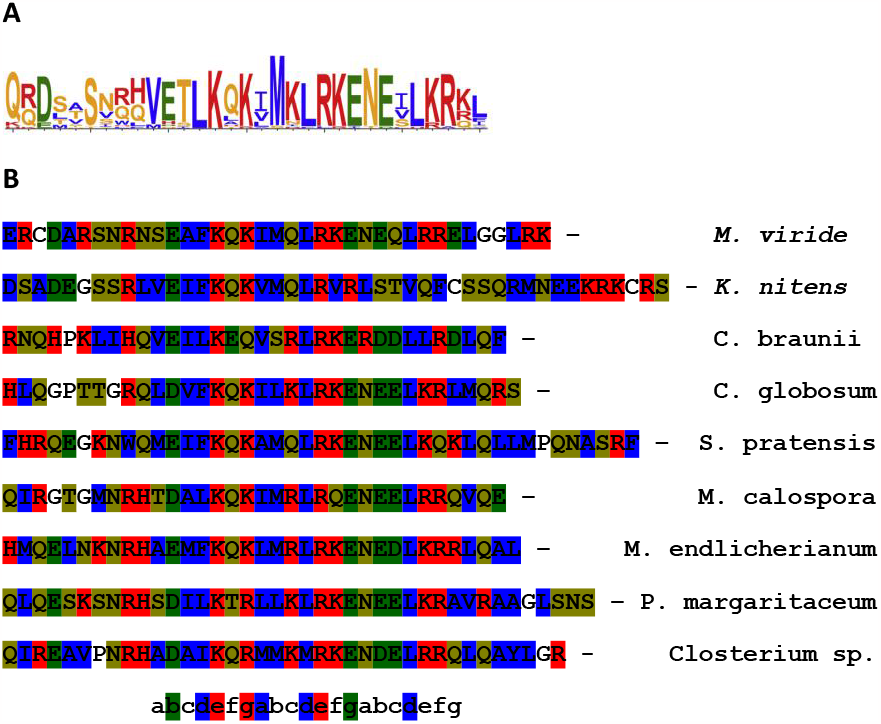
Comparison of the C-terminal sequences of the predicted 4/1-like proteins from charophyte alga. (A) Sequence logos of the C-terminal 4/1 conserved domains in flowering plants. Amino acids are colored according to chemical properties; polar (brown), hydrophobic (blue), negatively charged (green), positively charged (red). (B) Comparison of the C-terminal sequences of the 4/1 proteins from some charophyte alga. Positions of amino acid residues in the coiled-coil motifs are shown below the sequence alignment. *M. viride* - *Mesostigma viride* (class *[Mesostigmatophyceae](https://www.ncbi.nlm.nih.gov/Taxonomy/Browser/wwwtax.cgi?mode=Undef&id=96475&lvl=3&lin=f&keep=1&srchmode=1&unlock)*) (genomic sequence; NCBI accession - RPFO01000184); *K. nitens* - *Klebsormidium nitens (*class *[Klebsormidiophyceae](https://www.ncbi.nlm.nih.gov/Taxonomy/Browser/wwwtax.cgi?mode=Undef&id=131220&lvl=3&lin=f&keep=1&srchmode=1&unlock))* (genomic sequence; NCBI accession - BANV01000195); *C. braunii* - *Chara braunii (class [Charophyceae](https://www.ncbi.nlm.nih.gov/Taxonomy/Browser/wwwtax.cgi?mode=Undef&id=304574&lvl=3&lin=f&keep=1&srchmode=1&unlock))* (genomic sequence; NCBI accession - BFEA01000280); *C. globosum* - *Chaetosphaeridium globosum (class [Coleochaetophyceae](https://www.ncbi.nlm.nih.gov/Taxonomy/Browser/wwwtax.cgi?mode=Tree&id=304573&lvl=3&lin=f&keep=1&srchmode=1&unlock) )* (transcriptomic sequence; NCBI accession - JO169622); *S. pratensis* - *Spirogyra pratensis (class Zygnematophyceae)* (transcriptomic sequence; NCBI accession - GBSM01000070); *M. calospora* - *Mougeotiopsis calospora (class Zygnematophyceae)* (transcriptomic sequence; NCBI accession - GJZN01036048); *M. endlicherianum* - *Mesotaenium endlicherianum (class Zygnematophyceae)* (genomic sequence; NCBI accession - WELA01012513); *P. margaritaceum* - *Penium margaritaceum (class Zygnematophyceae)* (genomic sequence; NCBI accession - SZWC01002411); *Closterium* sp. - *(class Zygnematophyceae)* (genomic sequence; NCBI accession - CANIUD010000038).

The identification of protein regions with the land plant 4/1-like signatures provides an obvious support for further attempts to find the genes encoding the putative full-length 4/1 proteins in charophycean algae. Interestingly, many charophycean algae (except *Chlorokybus atmophyticus*, class *Chlorokybophyceae*) potentially encode proteins with C-terminal 4/1-like signatures that show strong conservation with land plant 4/1 proteins [10] (Figure 1). Indeed, most 4/1-like algal proteins contain a terminal coiled-coil element with an absolutely conserved Lys residue in the position ‘e’ and highly conserved Gln and Lys in positions ‘f’ and ‘g’ of the first heptad, respectively. Additionally, Arg and Lys are highly conserved in the positions ‘e’ and ‘f’, respectively, whereas E is highly conserved in the position ‘g’ of the second heptad. All three heptads contain conserved hydrophobic residues in the position ‘d’ (Figure 1). It has been speculated that this specific amino acid cluster may be involved in functionally important coiled-coil interactions with the other evolutionarily conserved plant protein(s) [4,5,10].

### 2.2. Prediction of full-length 4/1-like genes encoded by charophyte algae

After predicting of the 4/1 protein signatures, we used the gene finding program AUGUSTUS (http://bioinf.uni-greifswald.de/augustus/, accessed on 19 October 2023) in a way that provides automatic *ab initio* structural annotations of the putative full-length 4/1-like genes in charophycean genomes. The protein parts revealed by genomic TBLAST of the above genomic sequences (see section *2.1*.) and considered as peptide hints for AUGUSTUS were then further subjected to the following procedures: (i) genome segments encoding these peptides (15000 base pairs in the 5’ direction and 500 base pairs in the 3’ direction) were extracted, and (ii) further, the three reading frames in the positive strand of the extracted DNA segment were chosen to run AUGUSTUS using a probabilistic model. It should be noted that the length of the selected DNA segments is supported by data on the maximum 4/1 gene size in land plants (excluding gymnosperms) [10].

In our previous paper [12], we showed that the 4/1-like gene in the streptophyte alga *Chara braunii* (class *Charophyceae*) is intronless. However, our data in the present paper show that the predicted 4/1-like genes in other classes of charophycean algae contain multiple introns, the number which varies from four to seven (Figure 2). In particular, 4/1-like gene of the most basal streptophyte alga *Mesostigma viride* (class *[Mesostigmatophyceae](https://www.ncbi.nlm.nih.gov/Taxonomy/Browser/wwwtax.cgi?mode=Undef&id=96475&lvl=3&lin=f&keep=1&srchmode=1&unlock)*) [16] contains five introns, which is a lower number compared to 4/1 genes in angiosperms [10]. This fact can be explained either by true evolutionary process of gene origination or, by incorrect prediction of the exon positions. In the absence of publicly available assembled transcriptomic data for *M. viridae*, we performed primary and secondary structure comparisons of the predicted algal 4/1-like protein with those of land plants.

**Figure 2.**
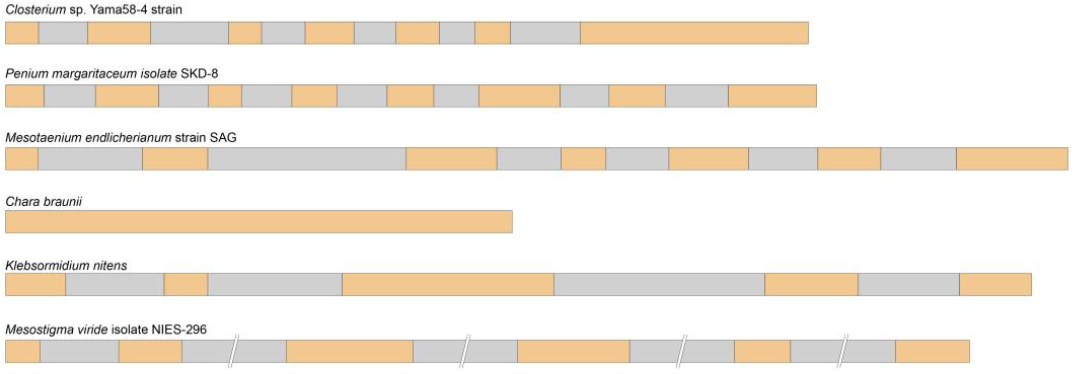
The exon/intron structure of the 4/1-like genes in representatives of different charophyte taxa. Exons are indicated by brown boxes, and introns are indicated by shaded boxes. Taxonomic positions for the presented algal species is shown on the text (section *2*). Total length of exons (amino acids) is shown in Supplementary figure 1. The representatives of class *Coleochaetophyceae* are not shown because of the lack of genomic sequence data.

Prediction of the protein secondary structure for the *M. viridae* 4/1 protein (http://www.compbio.dundee.ac.uk/jpred4/, accessed on 19 October 2023) reveals that this protein, like 4/1 proteins of land plants [4,10], is mostly alpha-helical with almost 85% residues involved in helices (Supplementary figure 1). Furthermore, direct sequence comparisons revealed 30% sequence identity of the whole algal protein with the *Nicotiana tomentosiformis* 4/1 protein (e-value – 6e-05) (Figure 3). These data strongly suggest significant accuracy in our prediction of the organization of the *M. viridae* 4/1-like protein. It should be noted that other predicted charophyte 4/1-like proteins also contain a high content of alpha-helices in their structure (Supplementary figure 1). In particular, amino acid residues in alpha-helices constitute 95% in *Klebsormidium nitens* protein, 93% in *Chara braunii protein, 92%* in *Mesotaenium endlicherianum*, 91% in *Penium margaritaceum* and 89% in *Closterium sp*. (Supplementary figure 1).

**Figure 3.**
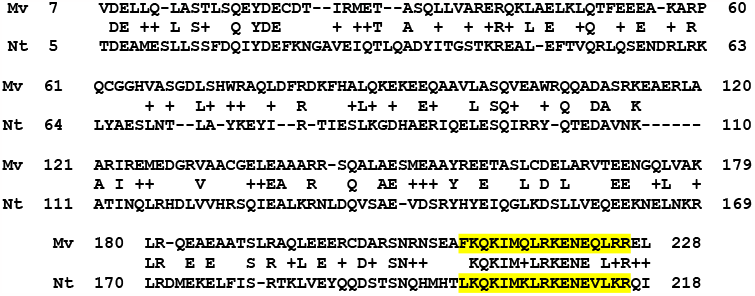
Pairwise sequence comparisons of the predicted amino acid sequence of algal 4/1-like protein from *Mesostigma viride* (Mv) and 4/1 protein of the selected flowering plant *Nicotiana tomentosiformis* (Nt). Numbers indicate amino acid position in the protein sequence. The C-terminal conserved coiled-coil signature is shown in the yellow.

### 2.3. Comparison of the 4/1 gene exon-intron organization in streptophyte algae and land plants

It is known that the exon/intron structure of single-copy genes can vary significantly depending on the taxonomic position of the plant species [21,22]. Our previous data for land plants and the present data for streptophyte algae confirm these observations (Table 1) [10] (Morozov et al., 2015). The 4/1 gene length of land plants ranges from 1142 bp in *Selaginella kraussiana* (lycophytes) to 213,214 bp in *Pinus taeda* (gymnosperms). This correlates with the variations in the intron size: in particular, intron length ranges from 47 bp to 64 bp in S. *kraussiana* and 103 to 109,876 bp in P. taeda [10].

**Table 1.** Gene size and intron length of the predicted charophyte 4/1 genes.

| Algae species | Total gene size/intron number | Largest and smallest introns |
| --- | --- | --- |
| *Mesostigma viride* (class *Mesostigmatophyceae*) | 6422 bp / 5 | 2811 bp – 129 bp |
| *Klebsormidium nitens* (class *Klebsormidiophyceae*) | 1698 bp / 4 | 349 bp – 163 bp |
| *Chara braunii* (class *Charophyceae*) | 834 bp / 0 | Not found |
| *Mesotaenium endlicherianum* (class *Zygnematophyceae*) | 1770 bp / 6 | 339 bp – 105 bp |
| *Penium margaritaceum* (class *Zygnematophyceae*) | 1350 bp / 7 | 105 bp – 75 bp |
| *Closterium* sp. (class *Zygnematophyceae*) | 1289 bp / 6 | 129 bp – 60 bp |

Similar correlations between single-copy gene and intron size and taxonomic position of plant species also have been found for the MSH1 gene, which is involved in the recombining and repairing of organelle genomes [22]. In general, it is known that the average intron size of gymnosperm species is much longer than that of other plants; for example, the average intron size of Chinese pine is 10,034 bp. In contrast, the average intron sizes of A. thaliana and O. sativa are 161 bp and 469 bp, respectively [22,23].

We show that the intron number of the predicted 4/1-like genes in streptophyte algae varies between species, ranging from 0 to 7 introns in C. brounii and *P. margaritaceum*, respectively. In contrast, the intron number in the land plant is relatively stable [10] (Morozov et al., 2015). In the class Zygnematophyceae of charophyte algae, the predicted 4/1-like genes possess rather short introns that are similar in size to those of lycophyte land plants. The largest introns are present in the *M. viride* 4/1 gene (Table 1) [10] (Morozov et al., 2015).

## 3. Discussion

This study provides an important breakthrough in the overall picture of the evolutionary history of the *4/1* gene in green plants (Viridiplantae). We extend the gene analysis of the *4/1* cistrons to six more algal streptophyte species in addition to multiple sequenced genomes of land plants [10]. Since *4/1-like genes* were not detected in species of *Chlorophyta, Rhodophyta* and *Glaucophyte* lineages, *it* is now clear that this gene originated in the key evolutionary events for Viridiplantae, namely, during the split of *Chlorophyta* and their descendants, streptophyte algae [24]. The phenotypes of the early-branching streptophytes (*Chlorokybophyceae* and *Mesostigmatophyceae*) are represented by simple one- or two-cell morphology. On the other hand, the *Klebsormidiophyceae* species, which diverged after *Chlorokybophyceae* and *Mesostigmatophyceae*, have evolved multicellularity by forming packets of cells [24]. Assuming our inability to find 4/1-like gene in species of class *Chlorokybophyceae*, it can be suggested that 4/1 gene first appeared in a single streptophyte algal progenitor 500–600 million years ago [25], and then it was lost by *Chlorokybophyceae* brunch. However, it was further evolved in *Mesostigmatophyceae* and *Klebsormidiophyceae* lineages [24].

In our previous paper, we have suggested that that intronless sequences of Chara 4/1 and other stress-related genes might have originated by retroduplication in ancestors of Charophyceae and undergone new intronization steps in early land plants [12]. However, new findings based on genome data for additional basal charophyte classes allow us to reconsider this view and show that 4/1 gene originated as intron-containing gene.

Experimental data and computer predictions of the secondary structure of 4/1 proteins in land plants have revealed that the 4/1 proteins consist of high proportion of alpha-helical regions [4,6]. Predictions made for 4/1-like proteins in charophyte algae also show significant percentage of helical residues. Many predicted alpha-helical regions represent coiled-coil (CC) domains (Supplementary figure 1) as it was also found for Nt-4/1 [6]. The high degree of similarity between the secondary structures of all 4/1 proteins suggests that these proteins have evolutionarily conserved amino acid clusters characteristic of myosins and kinesins, which could be involved in functionally important multivalent protein-protein interactions [4].

Despite extensive searches, no 4/1 genes have been identified in mosses, potato and tomato, although we assume that the ancestors of these plants must have possessed a 4/1 gene [4]. The absence of a 4/1 gene in the above mentioned species suggests that 4/1 proteins are probably not required for a vital primary metabolic function and functioning as accessory factors in stress response. It seems that there may be functional redundancy of proteins, and the lack of 4/1 genes in plants such as *Solanum* sp. and mosses may be compensated by other proteins [4]. The identification of proteins with the 4/1-like signatures in charophycean algae (see above) provides additional support for the hypothesis that the lack of 4/1 in mosses could be a result of gene loss in the immediate ancestor of Bryophyta.

## 4. Materials and Methods

Sequences for comparative analysis were retrieved from NCBI (http://www.ncbi.nlm.nih.gov/, accessed on 19 October 2023), and Phytozome (http://www.phytozome.net, accessed on 19 October 2023) databases. For computer-assisted secondary structure prediction, the JPred4 server (http://www.compbio.dundee.ac.uk/jpred4/, accessed on 19 October 2023) is used.

AUGUSTUS *ab initio* gene prediction was performed at (http://bioinf.uni-greifswald.de/augustus/, accessed on 19 October 2023).

Conserved sequence blocks in the Angiosperm 4/1 proteins were detected using WebLogo3 (http://weblogo.threeplusone.com/, accessed on 19 October 2023). The height of each column corresponds to the information content of the corresponding position in the alignment, and the size of the individual symbols within each column reflects the frequency of the corresponding amino acid at this position. The NCBI accession numbers for the annotated 4/1 sequences used for the sequence logo construction are as follows: *Populus trichocarpa* XM_002325222; *N. tabacum* EU117386; *Ricinus communis* XM002532486; *A. thaliana* NM_118735; *Arabidopsis lyrata* XM_002867538; *Oryza sativa* NM_001052428; *Sorghum bicolor* XM_002451499; *Zea mays* NM_001137007; *Hordeum vulgare* AK359619; *Brassica rapa* AC189360; *Capsicum annuum* GD066605; *Gossypium hirsutum* ES798773; *Malus domestica* GO550560; *Glycine max* FK638939; *Fragaria vesca* EX659494.

## Supplementary Materials

The following supporting information can be downloaded at: www.mdpi.com/xxx/s1, Figure S1: Predicted secondary structures in the selected 4/1 proteins from charophyte algae. Numbers indicate amino acid positions according the N terminus. Residues in secondary structure elements are denoted as H - alpha-helix, C – coiled coil structure.

## Author Contributions

Conceptualization, S.M. and A.S.; methodology, S.M.; software, A.S.; validation, S.M. and A.S.; formal analysis, A.S.; data curation, A.S.; writing—original draft preparation, S.M.; writing—review and editing, A.S. and S.M.; supervision, S.M.. All authors have read and agreed to the published version of the manuscript.

## Data Availability Statement

All data are available upon reasonable request.

