## Supplementary Figure 1 for "Occurrence of plant-specific 4/1 gene in streptophyte algae: brief view on the gene and protein evolution"

### *Mesostigma viride* isolate NIES-296 (*Mesostigmatophyceae*)

```
1---------11--------21--------31--------41--------51--------61--------71
MEAGKIVDELLQLASTLSQEYDECDTIRMETASQLLVARERQKLAELKLQTFEEEAKARPQCGGHVASGDLS
----HHHHHHHHH--------------HHHHHHHHH----HHHHHH-----HHHHHHHHH-----HHHHHHH
------------------------------------cccccccccccccc----------------------

--------81--------91--------101-------111-------121-------131-------141
HWRAQLDFRDKFHALQKEKEEQAAVLASQVEAWRQQADASRKEAERLAARIREMEDGRVAACGELEAAARR
HHHHHHHHHHHHHHHHHHHHHHHHHHHHHHHHHHHHHHHHHHHHHHHHHHHHHHHHHHHHHHHHHHHHHHH
---------------------------------------------------------------cccccccc

-------151-------161-------171-------181-------191-------201-------211
SQALAESMEAAYREETASLCDELARVTEENGQLVAKLRQEAEAATSLRAQLEEERCDARSNRNSEAFKQK
HHHHHHHHHHHHHHHHHHHHHHHHHHHHHHHHHHHHHHHHHHHHHHHHHHHHHHHHHHHHHHHHHHHHHH
cccccccccccccc-------------------CCCCCCCCCCCCCCCCCCCCCccc------CCCCCCC

-------221-------231
IMQLRKENEQLRRELGGLRK
HHHHHHHHHHHHHHHHH---
CCCCCCCCCCCCC-------
```

### *Klebsormidium nitens* (*Klebsormidiophyceae*)

```
1---------11--------21--------31--------41--------51--------61--------71
MPGLAVRTMTLLDEVMSTLEDVDVLQKEYNSVVDMNTKLKGSLMEHQMTNSLLEERCSELESERASLLEVHD
--------HHHHHHHHHHHHHHHHHHHHHHHHHHHHHHHH---HHHHHHHHHHHHHHHHHHHHHHHHHHHHH
------------------------------------------CCCCCCCCCCCCCCCCCCCCCCCCCCCC--

--------81--------91--------101-------111-------121-------131-------141
ESVRTALAQYDFREKYEKHLLSSADENDRQQKEAQRKTEALQTQLRAVREHESSERALRVAQIEKLQSEGS
HHHHHHHHHHHHHHHHHHHHHHHHHHHHHHHHHHHHHHHHHHHHHHHHHHHHHHHHHHHHHHHHHHHHHHH
-----------------------------------------------------------------------

-------151-------161-------171-------181-------191-------201-------211
KKDNAISALQQELEAAQRGLLEAKREKDEAQSKEQRKYQAELRLLQKTIKDMEESKLSLREEVKRAEHEA
HHHHHHHHHHHHHHHHHHHHHHHHHHHHHHHHHHHHHHHHHHHHHHHHHHHHHHHHHHHHHHHHHHHHHH
-----CCCCCCCCCCCCCCCCCCCCCCCCCCCC-cccccccCCCCCCCCCCCCCCCCCCCCCCCCCCCCc
```

```
-------221-------231-------241-------251-------261----
MTARMKFEEDSADEGSSRLVEIFKQKVMQLRVRLSTVQFCSSQRMNEEKRKCRS
HHHHHHHHHHHHHH-HHHHHHHHHHHHHHHHHHHHHHHHHHHHHHHHHHHH---
cccccc------------------------------------------------
```

### *Chara braunii* (*Charophyceae*)

```
1---------11--------21--------31--------41--------51--------61--------71
MLGEKKVDILEMTSELEREYKQTKAKCRELTAQLAREEVTRKCLEDRCQDLLRTNECLKQVNAECVAMAREQ
--------HHHHHHHHHHHHHHHHHHHHHHHHHHHHHHHHHHHHHHHHHHHHHHHHHHHHHHHHHHHHHHHH
------------cccccccCCCCCCCCCCCCCCCCCCCCCCCCCCCCCCCCCCCCCCCCCC-----------

--------81--------91--------101-------111-------121-------131-------141
YDFRKKSKELSIETEAKEDAWRRKERDWEKRVRDLNDELREERRQHKELKDKYLELSSRMEEEMRARRERE
HHHHHHHHHHHHH-HHHHHHHHHHHHHHHHH--HHHHHHHHHHHHHHHHHHHHHHHHHHHHHHHHHHHHHH
------------------------CCCCCCCCCCCCCCCCCCCCCCCCCCCCCCCCCCCCCCCCCC-----

-------151-------161-------171-------181-------191-------201-------211
KDWERQLRRLGDDMEDRRRRHEDLVASLEAQVAKTTDEIAVLRQENEAAAKRQQEEQLAADIKIAALLEK
HHHHHHHHHHHHHHHHH---HHHHHHHHHHHHHHHHHHHHHHHHHHHHHHHHHHHHHHHHHHHHHHHHHH
-----------------CCCCCCCCCCCCCCCCCCCCCCCCCCCCCCCCCCCcccccccccccCCCCCCC

-------221-------231-------241-------251-------261-------271-------
LQNADRRGSQLEGDLRKAEHTNTVIRTRYEEQLRNQHPRLIHQVEILKEQVSRLRKERDDLLRDLQF
HHHHHHHHHHHHHHHHHHHHH-HHHHHHHHHHHHHHHHHHHHHHHHHHHHHHHHHHHHHHHHHH---
cccccccccccccc----------------------cccccccCCCCCCCCCCCCCCCCCCCCC---
```

### *Mesotaenium endlicherianum* strain SAG (*Zygnematophyceae*)

```
1---------11--------21--------31--------41--------51--------61--------71
MTLTEVARQLLEDVDVLQQDYKEGKALVSNTKSLLHMEEVRRVSAEQHISSLDKVSFAFAFVYTVYHGRYND
--HHHHHHHHHHH--HHHHHHHHHHHHH----HHHHHHHHHHHHHH-------HHHHHHHHHHHHH---HHH
------------------------------------------------------------------------

--------81--------91--------101-------111-------121-------131-------141
VLHSLQYSYASKNQELMAQLELKDGQLKDVLKAHAMNMCNFTERAQAEKAHMNEKIRTTEKGLAEEKQQRL
HHHHHHHHHHHHHHHHHHHHHHHHHHHHHHHHHHHHHHHHHHHHHHHHHHHHHHHHHHHHHHHHHHHHHHH
--------------------------------------------------------------CCCCCCCCC
```

-------151-------161-------171-------181-------191-------201-------211

LAEQEMASLKRANESLKQKFDAAEATLMAKVGAENSEISLLRAKLHMEELEKQTAQEALQREKRHEAALT

HHHHHHHHHHHHHHHHHHHHHHHHHHHHHHHHHHHHHHHHHHHHHHHHHHHHHHHHHHHHHHHHHHHHHH

CCCCCCCCCCCCCCCCCCCCCCCCCC------cccccccccccccccccccccccccc------------

-------221-------231-------241-------251-------261-------

PPSYVARCTFSTAQEAKEKLEEHMQELNKNRHAEMFKQKLMRLRKENEDLKRRLQAL

HHHHHHHHHHHHHHHHHHHHHHHHHHHHHHHHHHHHHHHHHHHHHHHHHHHHHHH--

---------CCCCCCCCCCCCCCCCCCCCCccCCCCCCCCCCCCCCCCCCCCC----

### *Penium margaritaceum isolate SKD-8* (*Zygnematophyceae*)

1---------11--------21--------31--------41--------51--------61--------71

MTTSTCPDDSDILRAFDALQKEYFAGRAELMTVKSALAVSEAKRKSAELKASMQAKEIERLQQLRRADLQMT

---------HHHHHHHHHHHHHHHHHHHHHHHHHHHHHHHHHHHHHHHHHHHHHHHHHHHHHHHHHHH----

-------------------------------------------ccccccccccccccccccccc--------

--------81--------91--------101-------111-------121-------131-------141

KQHAHIAAEYEKLSLSYANDKQKWKKLHQDLEDSQLQRDASWKEVEQKLKERAENLEREFSEACSTIVSLE

HHHHHHHHHHHHHHHHHHHHHHHHHHHHHHHHHHHHH-----HHHHHHHHHHHHHHHHHHHHHHHHHHHHH

---------------------------------------------------CCCCCCCCCCCCCCCCCCCC

-------151-------161-------171-------181-------191-------201-------211

DKIASLSAEADNAREAARNGVGAVRQQYEKELEQLHAALHERERIIVELRNSLSQLKECYAASQLKLEEQ

HHHHHHHHHHHHHHHHHHHHHHHHHHHHHHHHHHHHHHHHHHHHHHHHHHHHHHHHHHHHHHHHHHHHHH

CCCCCCCCCCCCCCCC------------------------------------------------------

-------221-------231-------241--------252

LQESKSNRHSDILKTRLLKLRKENEELKRAVRAAGLSNS

HHHHHHHHHHHHHHHHHHHHHHHHHHHHHHHHHH-----

---------CCCCCCCCCCCCCCCCCCCCCCCCCCCC--

### *Closterium* sp. Yama58-4 strain (*Zygnematophyceae*)

1---------11--------21--------31--------41--------51--------61--------71

MLRESIIAVNGELRLVEIEIKKGQSELEVLRSALASAEYRREAAEQRELMLQNEVERLRSLRQAELEIIQQQ

--HHHHHHH---EEEEEEHHHH--HHHHHHHHHHHHHHHHHHHHHHHHHHHHHHHHHHHHHHHHHHHHHHHH

--------------------------CCCCCCCCCCCCCCCCCCCCCcccccccccccccc-----------

--------81--------91--------101-------111-------121-------131-------141

FHFRKLYEEAEDRLKKQQEEYDLKMEELKGVLSTVDDRASQAKTVAENRVRSLEKELSEAWAMVVTLEDRR

HHHHHHHHHHHHHHHHHHHHHHHHHHHHHHHHHHHHHHHHHHHHHHHHHHHHHHHHHHHHHHHHHHHHHHH

-------CCCCCCCCCCCCCCCCCCCCC----------------------ccccccccccccccccccccc

-------151-------161-------171-------181-------191-------201-------211

SLLITYVADESELATLRNQLHDEEARRKSLEKSLSMAEYKCKSPWPFSSLSWPVEYMGDSLTTVCAGSLA

HHHHHHHHHHHHHHHHHHHH-HHHHHHHHHHHHHHHHHHHH------EE---------HHHHHHHHHHHH

------------CCCCCCCCCCCCCCCCCCCCCccccccc------------------------------

-------221-------231-------241-------251-------261-----

ENQLQEQMKEFRQKMEEQIREAVPNRHADAIKQRMMKMRKENDELRRQLQAYLGR

HHHHHHHHHHHHHHHHHHHHHH------HHHHHHHHHHHHHHHHHHHHHHHH---

ccccccccccccccccccccc------CCCCCCCCCCCCCCCCCCCCCCCCCCCC
